# Side-Impact Collision: Mechanics of Obstacle Negotiation in Sidewinding Snakes

**DOI:** 10.1101/2020.04.22.055681

**Authors:** Henry C. Astley, Jennifer M. Rieser, Abdul Kaba, Veronica M. Paez, Ian Tomkinson, Joseph R. Mendelson, Daniel I. Goldman

**Affiliations:** U of Akron; Georgia Institute of Technology; Zoo Atlanta, Georgia Institute of Technology

## Abstract

Snakes excel at moving through cluttered environments, and heterogeneities can be used as propulsive contacts for snakes performing lateral undulation. However, sidewinding, often associated with sandy deserts, cuts a broad path through the environment that may increase the vulnerability to obstacles. Our prior work demonstrated that sidewinding can be represented as a pair of orthogonal body waves (vertical and horizontal) that can be independently modulated to achieve high maneuverability and incline ascent, suggesting that sidewinders may also use template modulations to negotiate obstacles. To test this hypothesis, we recorded overhead video of four sidewinder rattlesnakes *(Crotalus cerastes)* crossing a line of vertical pegs placed in the substrate. Snakes used three methods to traverse the obstacles: a Propagate Through behavior in which the lifted moving portion of the snake was deformed around the peg and dragged through as the snake continued sidewinding (115/160 runs), Reversal turns that reorient the snake entirely (35/160), or switching to Concertina locomotion (10/160). The Propagate-Through response was only used if the anterior-most region of static contact would propagate along a path anterior to the peg, or if a new region of static contact could be formed near the head to satisfy this condition; otherwise, snakes could only use Reversal Turns or switch to Concertina locomotion. Reversal Turns allowed the snake to re-orient and either escape without further peg contact or resorting to Propagate Through. We developed an algorithm to reproduce the Propagate Through behavior in a robotic model using a modulation of the two-wave template. This range of behavioral strategies provides sidewinders with a versatile range of options for effectively negotiating obstacles in their natural habitat, as well as provide insights into the design and control of robotic systems dealing with heterogeneous habitats.

## Introduction

Terrestrial habitats may have a variety of structural features with which moving animals must contend, including inclines, uneven terrain, granular substrates, and obstacles of various sizes and orientations. Obstacles are discrete structures in the environment, and thus animals may employ various behavioral strategies to traverse them, or may avoid them entirely (Collins et al., 2013; Daley et al., 2006; Kohlsdorf and Biewener, 2006; Li et al., 2015; Parker and McBrayer, 2016). However, traversing or avoiding obstacles may impose costs due to raising or deflecting the center of mass, altering limb kinematics, or force and energy needed to push itself through. Furthermore, improper obstacle negotiation can have serious consequences such as collisions, falls, and becoming stuck (Daley et al., 2006; Li et al., 2015; Parker and McBrayer, 2016).

Snakes present an intriguing counterexample to the obstacle-negotiation issues in limbed taxa. Unlike limbed organisms, which typically switch between locomotor modes in response to changes in speed (e.g. walking, running, and galloping) (Hildebrand, 1985), snakes may change locomotor mode in response to habitat (Gans, 1975; Gray and Lissmann, 1950; Jayne, 1986). Additionally, the most common locomotor mode in snakes, lateral undulation, uses the same discrete environmental structures, which are obstacles to limbed animals, as push points for generating propulsive force (Gray and Lissmann, 1950). As a consequence, while obstacles typically slow down limbed animals(Clifton et al., 2020; Collins et al., 2013; Hyams et al., 2012; Sponberg and Full, 2008), the presence of these structures allows snakes to switch locomotor modes from concertina (generally slow (Jayne, 1986)) to lateral undulation (the fastest mode (Jayne, 1986))(Astley and Jayne, 2009). As such, snake speed during lateral undulation increases with increasing obstacle density (except at extreme densities in which snakes no longer have room to use lateral undulation effectively) (Kelley et al., 1997).

Obstacles typically pose few challenges to non-sidewinding snakes, as a minor reorientation of the head will lead the body around the obstacle, either actively or due to passive collision with the obstacle itself (Schiebel et al., 2019). However, sidewinding may be uniquely susceptible to disruption by obstacles, due to the broad path the body takes through the environment (Gray, 1946; Jayne, 1986; Mosauer, 1930; Mosauer, 1932a; Mosauer, 1935)(Fig. 1). This mode of locomotion, consisting of a simultaneous superimposed vertical and lateral undulations resulting in propagating regions of static contact and lifted movement (Astley et al., 2015; Marvi et al., 2014), is the primary locomotor mode in a few species of desert-dwelling vipers (Brain, 1960; Gans and Mendelssohn, 1972; Gray, 1946; Mosauer, 1932b), but is seen more widely in particular environmental conditions (Jayne, 1986; Tingle, 2020). In all cases, the static contact of lowered segments of the body allows the snakes to gain traction without significant substrate yielding (Marvi et al., 2014). These habitats also include few obstacles to disrupt the broad path of sidewinding snakes (Fig. 1A) (Cowles, 1956; Mosauer, 1935). But what are the consequences of a sidewinding snake which cannot or does not avoid an obstacle in its path?

**Figure 1.**
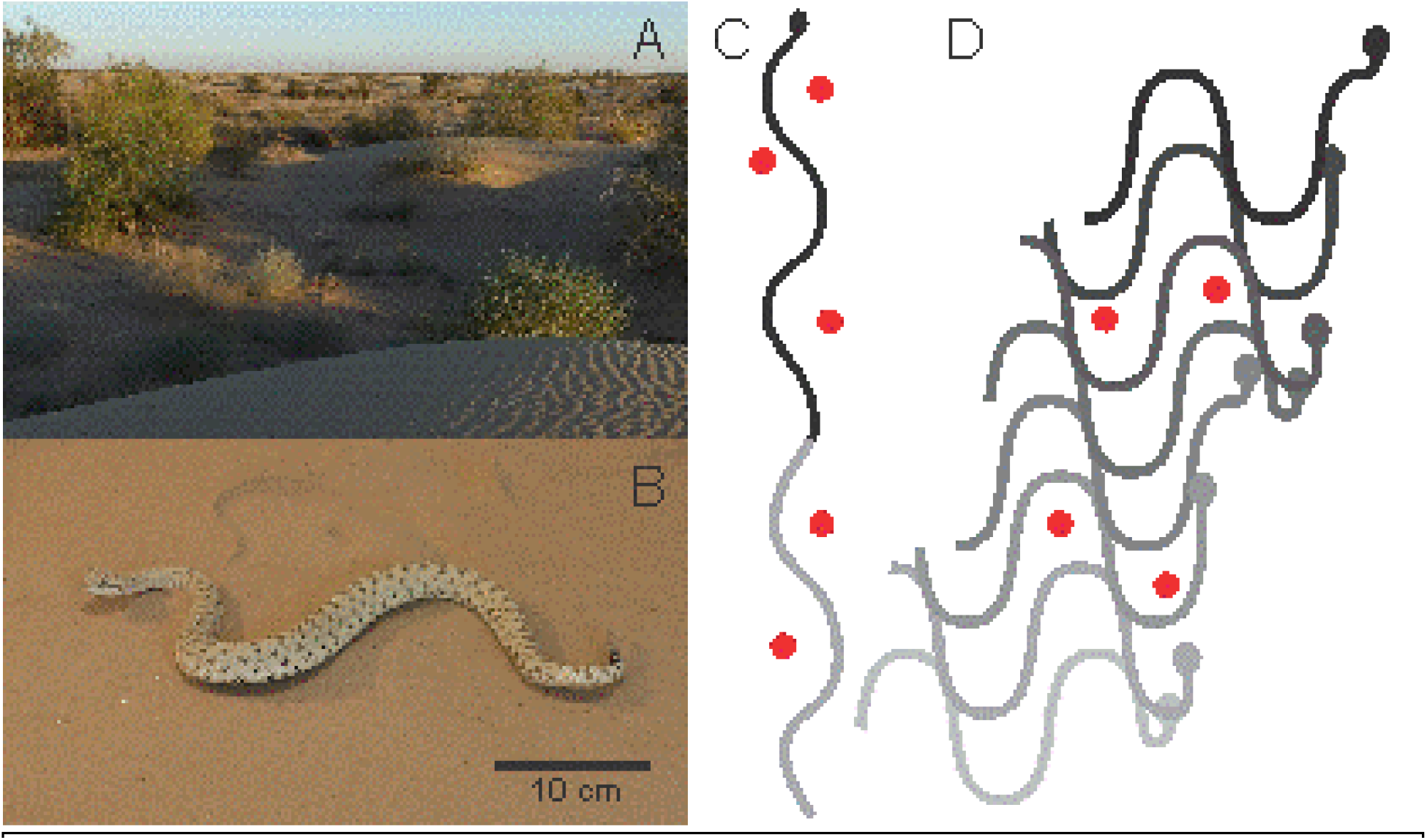
Snake Locomotion and Obstacles. A) Typical habitat for sidewinder rattlesnakes *(Crotalus cerastes)* (Photo by HCA). B) A sidewinder rattlesnake sidewinding on sand (Photo by HCA). C) A diagram of lateral undulation without slipping, with obstacles in red, showing the narrow path through the environment which allows total evasion of the obstacles. D) A diagram of sidewinding without slipping, with obstacles in red, showing the broad path the body takes through the environment and consequent necessity of obstacle interaction. C & D based on Gray 1946.

Sidewinding can be represented as a combination of offset waves of vertical and horizontal undulation on the body, and applying this control template to a robotic snake will successfully produce sidewinding (Burdick et al., 1993; Jayne, 1986; Jayne, 1988). Our prior work showed that each wave could be modulated independently to ascend inclines (Marvi et al., 2014) and maneuver (Astley et al., 2015), suggesting that further modulations could be the basis for any observed obstacle negotiation. Alternatively, snakes may simply abandon sidewinding when confronted by an obstacle. If multiple strategies can be used, what determines the selection of a strategy, and what are the performance consequences of each? We examined the behaviors and kinematics of sidewinder rattlesnakes (*Crotalus cerastes*) via overhead video as they traversed an array of vertical pegs embedded in a natural-sand substrate to examine the behavioral strategies used, as well as their and consequences and circumstances, followed by testing a template-based modulation strategy in a robophysical model..

## Materials & Methods

### Animals

We studied obstacle negotiation in four adult sidewinder rattlesnakes (*Crotalus cerastes*) (mean ± s.d.: total length 47.1 ± 5.0 cm, mass 114.2 ± 41.7 grams) collected near Yuma, Arizona, USA, and housed at Zoo Atlanta. All trials were conducted more than three days after feeding and before any visible signs of the molt cycle. Trials were conducted at temperatures from 21–25° C, comparable to the body temperatures of snakes collected in the field (Brattstrom, 1965). Experimental protocols were approved by both Georgia Institute of Technology Institutional Animal Care and Use Committee and Zoo Atlanta Scientific Review Committee.

### Obstacle Trials

Locomotion trials were conducted in a 1 x 2 m^2^ arena filled with sand from the snakes’ capture locality to a depth of approximately 7 cm. This arena was a fluidized bed (Marvi et al., 2014), capable of forcing air through a porous sheet beneath the sand to fluidize the sand and return it to a standard starting state while erasing tracks. All trials were recorded for their full duration using an overhead 1080p webcam (Logitech C920 HD Pro Webcam) in a fixed position, calibrated using images of a meter stick.

Preliminary experiments were conducted with three conditions to determine the effect of a variety of obstacles: a series of ridges, hemispherical obstacles, and narrow vertical cylinders (henceforth “pegs”). Five sand ridges running across the short axis of the arena were constructed by moving the sand manually, with each ridge being approximately 10 cm tall and 21 cm across the base, with both sloped sides at the maximum slope possible for this sand (28°). Snakes were encouraged to cross these ridges along the long axis of the arena, and did so with no apparent difficulty or reluctance, nor any obvious modifications to waveform beyond raising the head to look above the peak of each ridge and consequently moving a greater distance on that cycle; this experimental treatment was discontinued after preliminary trials (Sup. Vid. 1). Hemispherical obstacles, 100 mm diameter, were 3D-printed using ABS plastic. Each had a flat protrusion in a “+” shape on the flat surface to anchor the obstacles into the sand. Snakes often attempted to avoid the hemispheres but, upon encountering them, the snakes were able to lift the moving regions of their body higher and drag them across the surface of the hemisphere with minimal loss in speed and no apparent difficulty (Sup. Vid. 2). Finally, vertical pegs (3D printed ABS) of 6.35 mm diameter and 15.4 cm height were implanted into the sand. Pegs were anchored beneath the sand via approximately 15 x 15 cm sections of 2.54 cm thick aluminum honeycomb (Part No. 9635K5, McMaster Carr), with a 3 mm diameter 2.54 cm long protrusion from the bottom of the peg inserted into one of the cells of the honeycomb. The use of a honeycomb structure allows the sand to be fluidized with the obstacles in place, although the obstacles had to be held in place during fluidization. Approximately 10 cm of peg was protruding vertically above the sand during trials (~4 times body height). The vertical pegs were the only obstacle to prompt noticeable modulations of sidewinding, and were used in all experimental trials presented in this paper.

### Vertical peg trials

Vertical pegs were arranged in a 1-meter row across the short axis of the arena at intervals of 10 or 15 cm (approximately 4- and 6-times body width), dividing the arena into two 1×1 m^2^ areas. Preliminary trials with peg rows oriented along the long axis of the bed did not allow sufficient preparatory area for the snakes and snakes frequently collided with the arena wall before completing transit of the peg row. Before each trial, peg anchors were buried in the sand by hand, then held in place with a meter stick while the sand was fluidized. Fluidization was performed between “trials”, defined as a series of successive movement bouts (“runs”) of the same snake between rest periods, but the bed was not fluidized between runs due to the large number of runs required. Snakes did not perceptibly alter their behavior when crossing disturbed sand. Trial duration was limited to 15 minutes, and snakes were given a minimum of 10 minutes of rest between trials; trials were terminated for the day if snakes showed signs of fatigue (rapid increase in cycle duration, stopping upon any peg contact, refusal to move even when contacted with a snake hook, adopting defensive postures and behavior). Snakes were encouraged to move in the direction of the pegs via body movements of the experimenters, but were not physically contacted with any objects at any point during a run.

We performed 38 trials (N = 23 at 10 cm spacing, N = 15 at 15 cm spacing) with an average of 26.3 runs per trial for a total of 998 runs. Some runs (42.7% of the total) were discarded or halted due to the snakes moving in a direction which would not intersect the peg line, while in an additional 9.2% of runs, snakes turned to avoid the peg line (7.6% differential turns, 1.6% reversals (Astley et al., 2015))). Additionally, 32.1% of runs were excluded because the snake came to a complete stop during the run (unless followed by concertina locomotion within one second) or if the snake contacted or interacted with the arena wall before, during, or within half a cycle after contact with one or more pegs. The final data set included 16% (N = 160) of runs for analysis. Analyzed runs were roughly evenly split (84:76) between the two possible vertical phase offsets corresponding to “head on the right” and “head on the left” (Astley et al., 2015; Marvi et al., 2014).

### Analyses

Runs were categorized into three broad categories based on the behavior of snake when traversing the pegs: “Propagate Through”, in which the snake continues sidewinding while deforming the body to move past the pegs (Sup. Vids. 3–6), “Reversal”, in which the snake performs a reversal turn (Astley et al., 2015) upon contact to either approach the peg line from a new direction or to evade any other peg contact entirely (Sup. Vid. 7, 8), and “Concertina” in which the snake stopped sidewinding and instead used one or more cycles of flat-surface concertina locomotion (Gans, 1970; Jayne, 1986) to move past the peg line (Sup. Vid. 9). An additional classification was used to subdivide the runs more finely, denoting cases when snakes utilized two behaviors in sequence, such as a Reversal followed by Propagate Through. In all such cases, the first behavior was used as the broad categorization.

Videos of runs were loaded into ImageJ (Schneider et al., 2012), and a video frame was selected. Digitized points were placed along the snake’s midline using the “segmented line” tool, and a custom macro-program was used to fit a curved spline to these points and interpolate 128 evenly spaced points along the body from the tip of the nose to the base of the rattle, the coordinates of which were saved. For each run, frames were selected at the start and end of contact with the peg line, as well as at times at approximately one half or one cycle of sidewinding before and after contact, for a total of six frames. In “Propagate Through” responses, an additional frame was digitized midway between the start and end of peg contact. Points were also digitized for each peg which was contacted by the snake, in order from anterior to posterior, though pegs beyond the first proved uninformative, as snakes contacted more than one peg only 35/160 runs with the second contact usually on the last half of the body, and a third peg contact occurred only twice.

These midlines were analyzed using a custom script in MATLAB (2015a, Mathworks Inc., Natick, MA, USA) to generate variables after up-sampling the 128 body points to 500 points using a spline interpolation. The location of the centroid of the body was the mean position of all 500 points, which was used to compute speed, overall direction, and change in direction for the three successive frames leading up to and including initial contact and the three successive frames including and following final peg contact. To determine whether traversing the peg line adversely affected the snakes, we computed the relative speed of traversal, defined as the average speed during the period from first and last peg contact divided by average speed prior to peg contact. The orientation of the velocity vector, relative to a vector from the body centroid to the head, was used to determine whether the snake was sidewinding with its head on the left or right, each of which corresponds to a vertical-wave phase shift relative of ± π/2 relative to the horizontal wave (Astley et al., 2015).

Local curvature (inverse of radius-of-curvature) was computed along the spline by determining the radius of a circle that intersected three points (the point under consideration and two additional points, each 30 points (6% of body length) anterior or posterior to the first; as a consequence, curvature could not be calculated for the first and last 6% of the body. The sign of the cross-product of vectors from the posterior point to the current point and the current point to the anterior point was used to determine whether curvature was to the left or right, and left curvature was given a negative sign. Curvature changed cyclically along the length of the body, and the points closest to zero curvature (straight) were used to determine half-wavelength. Local curvature allowed computation of horizontal wave phase (Astley et al., 2015), which, combined with knowledge of the relative phase offset between vertical and horizontal waves based on head position relative to velocity vector, allowed segments to be designated as moving or static (Astley et al., 2015). This allowed calculation of the total number of static contact regions and the angle of tracks left by the sidewinder (a measure of body posture). The track angle was determined by selecting the point of zero curvature on the body, at which point the body should be parallel to the tracks (Fig. 1) and calculating the angle between the body and the forward velocity vector. Half wavelength, number of static contacts, and track angle were calculated for the snake before contact, at initial peg contact, at the end of peg contact, and after contact, with an additional half-wavelength calculation midway through traversal in “Propagate Through” responses. Because less than a complete cycle occurred between successively digitized frames before and after peg contact, the phase difference for each point in two successive frames, averaged across the body length, and time elapsed between frames can be used to calculate cycle frequency before and after contact.

To test if the location and nature of the peg contact on the snake’s body determined the categorical response, we calculated several variables related to peg contact. For each peg contact, we recorded the position along the body from head to tail and the phase and curvature at the point of contact, as well as recording the number of pegs contacted. Further analysis focused on the role of the most anterior peg contact; almost all second peg contacts occurred on the posterior half of the body which, if the site of the only contact, nearly always produced a Propagate Through reaction, and third contacts were too rare to analyze. Of particular interest was the relationship between the first peg contact and the static region of the body. Observations of several Propagate Through runs suggested that sidewinders were primarily pulling the body past the pegs and through the gap between the peg and the first static contact, as evidenced by the “slipping” of more posterior body points along the ground during these runs, including one run in which the entire snake slipped in the sand with the apparent force applied to the peg (Sup. Vid. 5). These suggested that the snake required a static contact point anterior to the peg contact location, which would serve as an anchor point from which the snake pulled the posterior portion of the body forward during the Propagate Through response. To determine the effect on behavior, we extrapolated a line from a linear regression of the anterior-most static points of low horizontal body curvature (less than half the maximum curvature at all static points) to represent the predicted future propagation direction of the static region, and calculated the orthogonal distance of the peg from this line (Fig. 2A). This variable was termed the “orthogonal distance to static contact”, and negative values denoted cases where the propagating static region would pass posterior to the peg and leave the snake without static contacts anterior to the peg from which to pull the body forward, while positive values indicated the static region would propagate anterior to the peg and allow static anterior anchor points (Fig. 2A). This variable was expressed as both a distance and a simple categorical variable for whether the peg was anterior or posterior to the predicted propagation line. Finally, we noted whether a new anterior static contact point was formed after initial peg contact in cases of Propagate Through only (Fig. 2BC), as this variable could not be meaningfully evaluated in the other behaviors.

**Figure 2.**
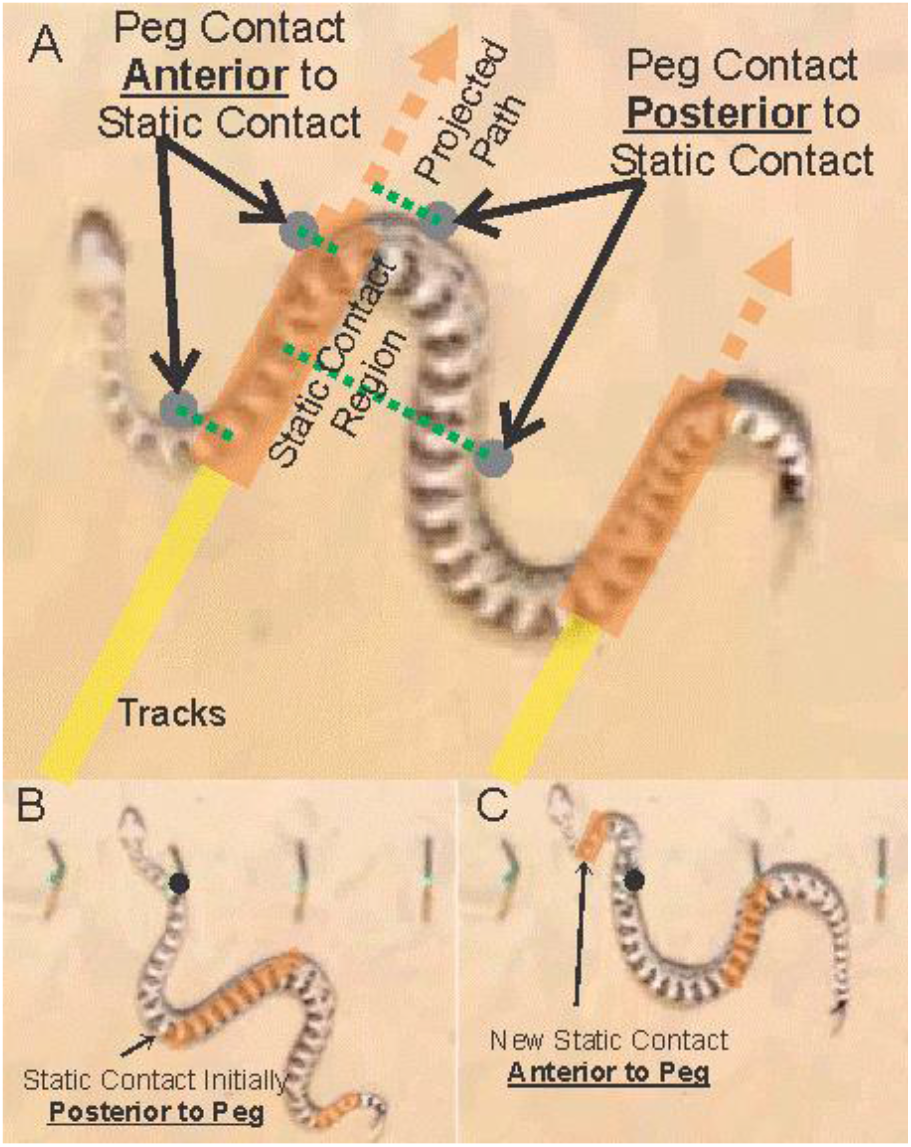
Contact locations relative to static regions. A) A sidewinder rattlesnake in motion, showing four potential contact locations (black circles). Orange shaded regions are portions of the body in static contact with the ground, yellow lines are tracks, and dashed orange lines represent projected future static contact locations. Green dashed lines show the orthogonal distances of each contact location to the static contact region; those considered anterior and posterior to the anterior-most static contact region are labeled. B) A snake makes contact with a peg during a run, with the peg being categorized as anterior to the static contact region. C) In a subsequent frame of the same video, the snake establishes a new static contact region anterior to the peg, and thus converts the peg contact categorization from anterior to posterior to the static contact region.

### Robophysical Model

To test whether modulations of the two-wave template could reproduce the Propagate Through behavior, we used a robophysical model in a dedicated test arena. From our biological experiments, we observe that the strategy employed by animals to move beyond rigid obstacles includes (1) an increase in amplitude that is initiated near the head of the animal and passed down the body and (2) the creation of a static contact with the substrate near the head of the animal. To test the importance of these observations in successful obstacle negotiation, we created a sidewinding robot from 15 Dynamixel AX-12A servo motors (Fig. 3A). Motors were linked together with custom 3D-printed brackets, and the direction of actuation alternated down the body such that the angular positions of odd-numbered motors varied in the horizontal plane and even-numbered motors were actuated in the vertical plane (Wright et al., 2012), with a total of 8 horizontal motors and 7 vertical motors. Using our previously discovered two-wave template (Astley et al., 2015; Marvi et al., 2014), each set of motors was programmed such that angular positions varied sinusoidally down the body, such that

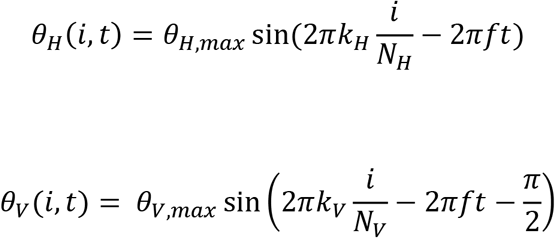

**Figure 3.**
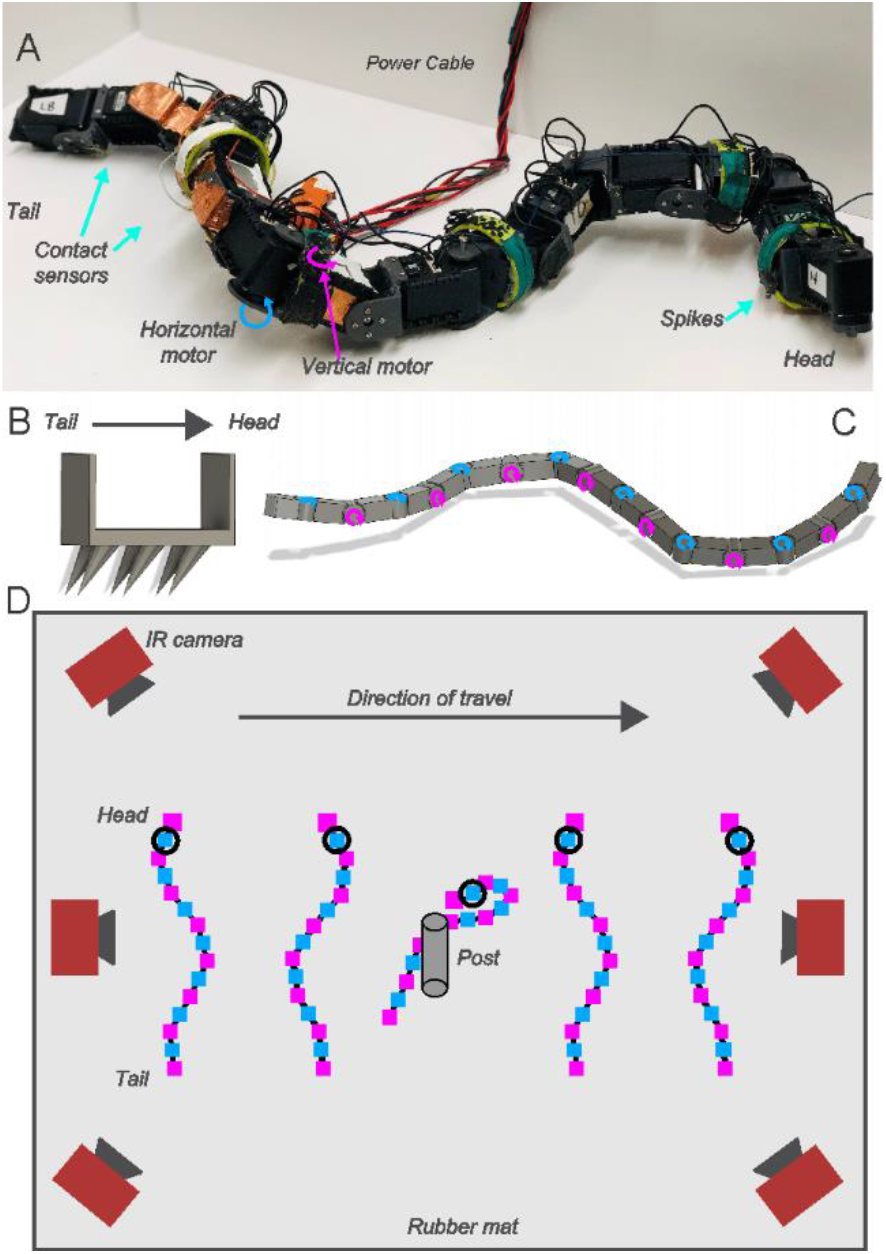
Robot obstacle negotiation. (A) A photograph of a sidewinding robot. Sixteen Dynamixel AX-12A servo motors were linked together with 3D printed brackets. The orientation of the motors alternated down the body, such that the direction of actuation of odd motors along the body is in the horizontal plane and the angular position of even motors varies in the vertical plane. Contact sensing is achieved through capacitive touch sensors affixed to the lower half of the body. (B) 3D printed angled spikes were affixed to the bottom of the second motor to reduce slipping of the robot’s contact point with the substrate. (C) Schematic of the snake robot, showing alternating left and right motors and directions of actuation (D) Schematic of the arena in which experiments were performed. Six IR-sensitive cameras captured the positions of eight reflective markers as the robot moved within the arena. A single vertical post was rigidly affixed to the rubber substrate. Upon establishing contact with the post, the robot either continued its nominal waveform or increased angular amplitude to pull past the post.

Where subscripts Hand *V* refer to, respectively, horizontally and vertically oriented motors; *θ* is angular position of a specified motor, *i*, at time, *t. θ_max_* is the amplitude of the sinusoidally-varying angular position of each motor, *k* is the number of waves along the body, *N* is the number of motors, and *f* is the temporal frequency of oscillation. Parameters used for experiments presented here are shown in the table below.

| | $\theta_{max}$ [rad] | $\Delta\theta_{max}$ [rad] | $k$ | $N$ | $f$ [Hz] |
| --- | --- | --- | --- | --- | --- |
| $H$ | 1.05 | 1.5 | 1.5 | 8 | 0.26 |
| $V$ | 0.225 | 0 | 1.5 | 8 | 0.26 |

We added contact sensing to the side of the back half of the robot to test different animal-inspired control strategies for successfully moving beyond the obstacle. Contact sensing consisted of 8 copper sheets on the 8 most posterior segments, connected to an MPR121 capacitive touch-sensor breakout board, that interfaced with the robot controller via I2C communications. To test the efficacy of increasing the horizontal amplitude, we programmed the robot to increase its horizontal amplitude for one cycle, *θ_H,max_ → θ_H,max_* + Δ*θ_H,max_*. To ensure continuous and smooth robot motion, the change in amplitude was initiated at the head at the beginning of an undulation cycle (so *θ* = 0) and subsequently passed to horizontal motors along the body for one cycle, after which the amplitude was returned to its nominal value. To test the effect of the anchor point near the head, we added 3D-printed spikes to the ventral side of the second motor. The spikes (Fig 3B) engaged with the underlying rubber substrate and reduced sliding motion. An infrared-reflective marker was added atop each horizontal motor, and six Natural Point Optitrack Flex-13 cameras recorded and tracked the position of each marker through time at 120 Hz as the robot moved within a 1.2 m x 1.2 m arena (Fig 3C). A single vertical post was rigidly affixed to the underlying substrate. To test how sensitive the system is to contact location and body configuration at the time of contact, we examined a range of starting positions for the robot by placing the tail at one of 16 starting points, arranged in a 4×4 grid. The long axis of the robot was parallel in all starting configurations, but these slight differences resulted in contact occurring at different locations on the body and times during the cycle. A total of 3–5 trials were performed for each starting position and behavior.

### Statistical Analyses

All statistical analyses were performed in JMP Pro v12 (SAS Institute, Cary, NC, USA). For comparisons between pre- and post-obstacle negotiation of variables and the effect of peg spacing, two-tailed t-tests and Pearson’s χ^2^ tests were used, while for comparisons among response types, a one-way ANOVA was used. Individual could not be considered as a random factor in a repeated measures ANOVA because the not all individuals had three or more Concertina responses.

To determine which variables predicted the behavior used to negotiate the obstacles, we performed a series of logistic regressions, with the broad behavior category (Reversal, Concertina, Propagate Through) as the dependent variable. Independent variables were the static phase, cycle frequency, speed, turning, trajectory angle, and half wavelength prior to peg contact, the number of static regions and number of pegs contacted at the time of peg contact, and the curvature, phase, location along the body and orthogonal distance to static contact of the anterior-most peg contact. Variables were ranked according to Akaike Information Criterion (with correction, AICc), considering only variables with a *χ*^2^ significant at p < 0.05. Models were compared by calculating relative likelihood, which shows the probability that a given model will have lower information loss than the one with the lowest AICc.

For robot trials, we defined the start of the interaction identifying the first time at which the established contact with the post (determined from tracking data, when the smallest distance between the post center and the robot marker was less than robot width plus post diameter, 7.7 cm). The robot was considered “stuck” or “pinned” by the post until the minimum distance between the robot and the post exceeded 20 cm. This number was chosen to be larger than the initial contact threshold because the robot slid around and would temporarily break contact with the post before re-establishing contact and continuing to spin around the post. To compare performance across initial conditions and behaviors, we normalized the pinning time, *t_pin_* by the total experiment time, *t_exp_*. *t_pin_/t_exp_* = 1 indicated that the robot remained “pinned” to the post throughout the entire experiment, while anything less than unity indicates that the robot successfully moved beyond the post. Performance for each behavior was determined based on the cumulative density function (CDF) of *t_pin_/t_exp_*.

## Results

### Snake Behavioral Responses to Obstacles

The responses of sidewinders to impact with obstacles could be broadly grouped into three categories, termed “Propagate Through”, “Reversal” and “Concertina”. Propagate Through was the most common (71.9%, 115/160) broad behavioral response, and was characterized by a continuation of sidewinding locomotion with no vertical-wave phase changes and modest directional changes (Fig. 4). In Propagate Through responses, the lifted, moving portion of the body deforms as it is moved past the peg, causing slipping at posterior static contact points but not anterior ones (Fig. 4, Sup. Vid. 3). Reversals are a type of turning behavior caused by a 180° phase shift in the vertical wave, in which the lifted, moving body segments and static, ground-contact segments switch roles, resulting in an abrupt change in direction (Astley et al., 2015) (Fig. 5, Sup. Vid. 7). Reversals were initiated upon contact with the peg in 35/160 (21.9%) runs. In the remaining 10 runs (6.3%), sidewinders stopped performing sidewinding upon contact with the obstacle and instead performed one or more cycles of flat-surface Concertina locomotion (Fig. 6, Sup. Vid. 9)(Gans, 1970; Jayne, 1986). Flat-surface concertina locomotion is similar to tunnel and arboreal concertina locomotion, except that instead of anchoring against the substrate by exerting lateral force on the tunnel walls (Jayne, 1986) or a medial grip on the branch (Astley and Jayne, 2007), the static portion provides anchoring only by static friction with the substrate (Gans, 1970; Jayne, 1986). During flat-surface concertina locomotion, the anterior portion of the body is flexed into tight horizontal waves that maintain static contact with the substrate as the posterior body is straightened and pulled forwards, after which the anterior portion is straightened and pushed forwards as the flexed posterior body waves serves as a static anchor (Fig. 6). Unlike sidewinding (in which body points also show cyclic periods of stasis and movement), in concertina locomotion the body waves do not propagate posteriorly, and are straightened and reformed with each cycle, though regions of movement and stasis do propagate (Fig. 6)(Astley, 2018; Jayne, 1986).

**Figure 4.**
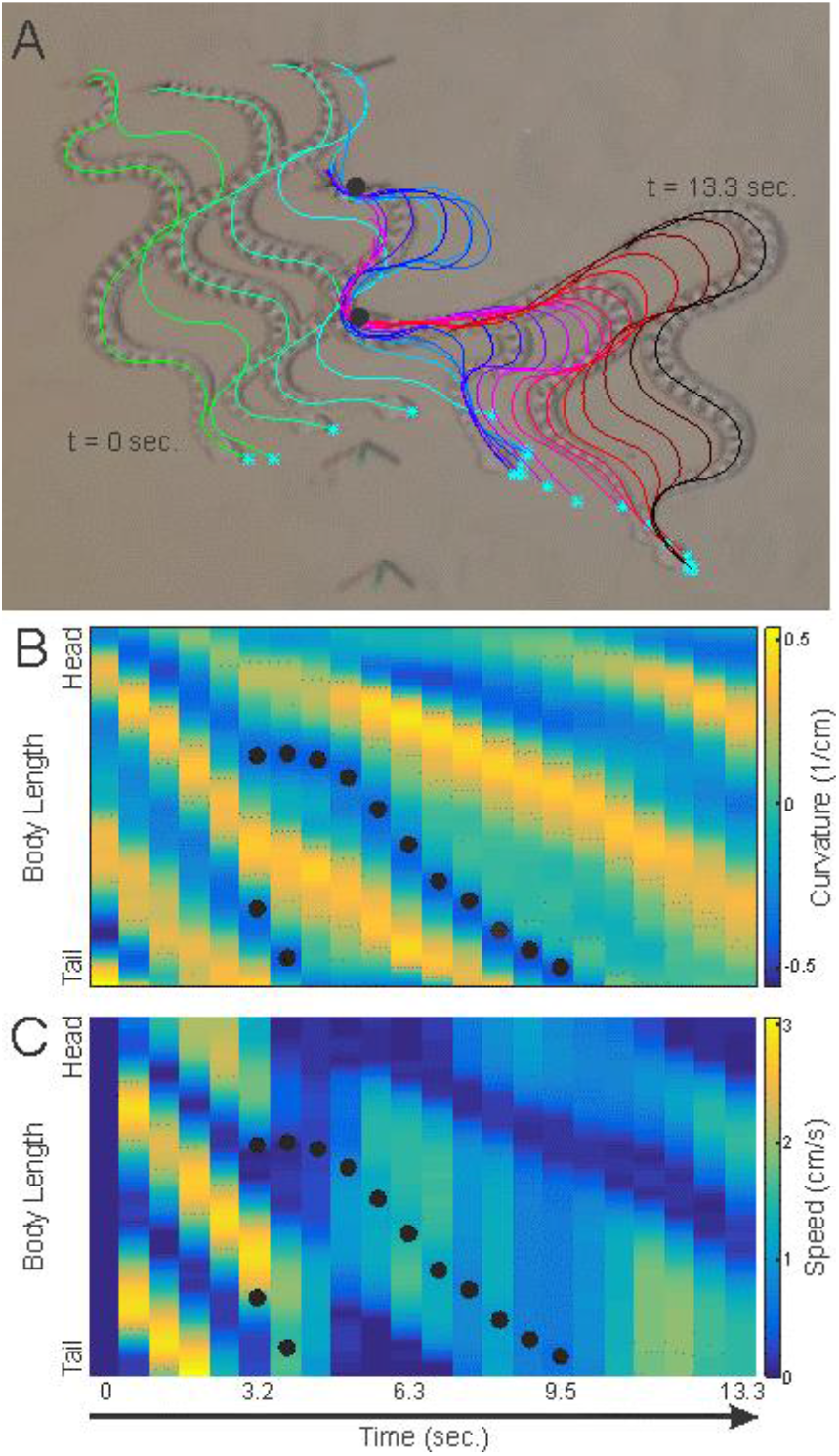
Propagate Through response. A) Sequential images from Sup. Vid. 3 showing a Propagate Through response, with midline traces at 0.63 second intervals showing the path of the body. The snake is moving from left to right. B) Curvature and C) Speed (derived from midline traces) along the snake’s body over time. The vertical axis is along the length of the body, while the horizontal axis is over time. Bands of curvature and speed going from top left to bottom right show posterior propagation of body waves and moving/static segments. Black circles indicate the location of the peg contact along the body for sequential frames.

**Figure 5.**
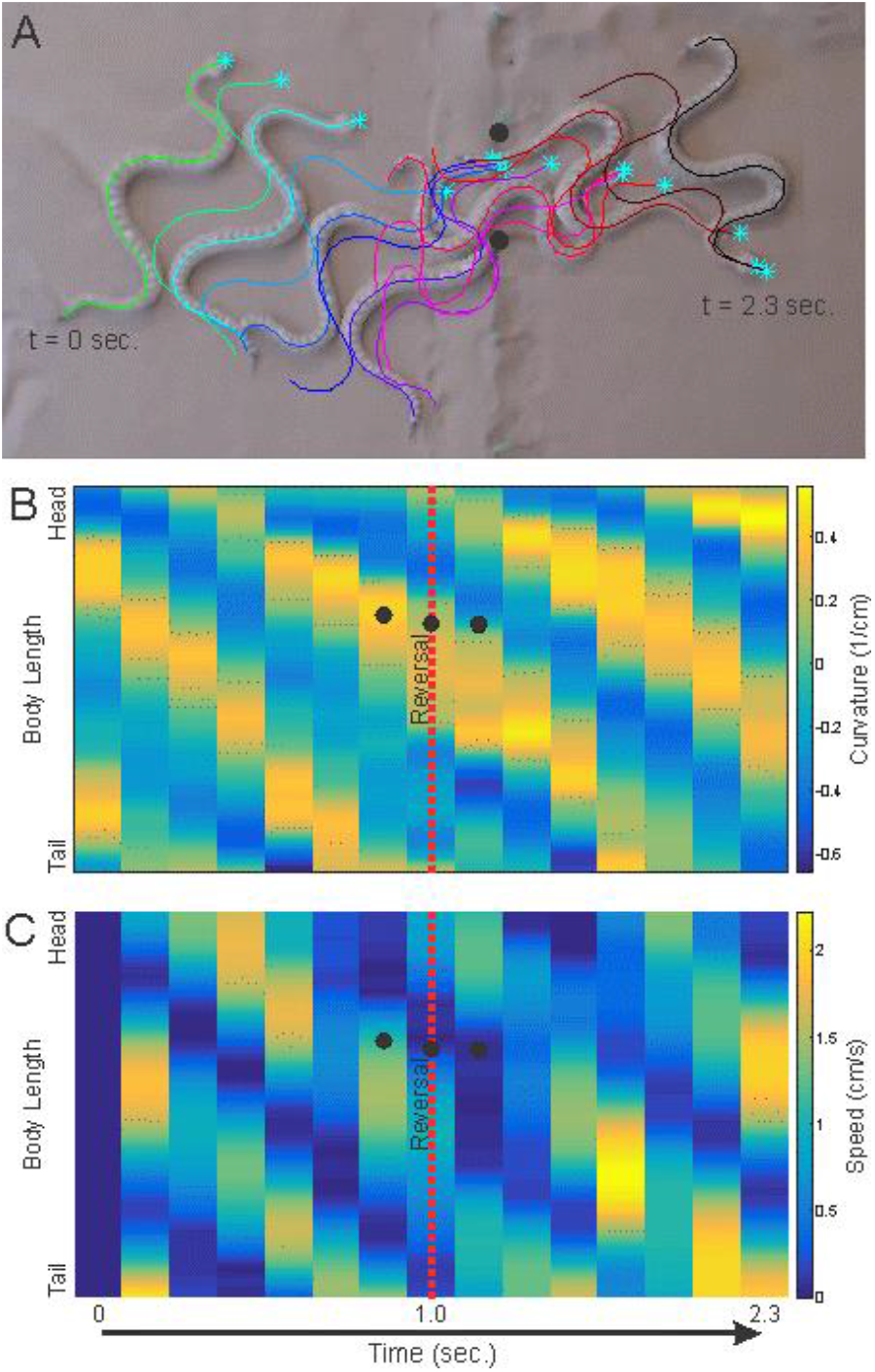
Reversal response. A) Sequential images from Sup. Vid. 7 showing a Reversal response, with midline traces at 0.17 second intervals showing the path of the body. The snake is moving from left to right. B) Curvature and C) Speed (derived from midline traces) along the snake’s body over time. The vertical axis is along the length of the body, while the horizontal axis is over time. Bands of curvature and speed going from top left to bottom right show posterior propagation of body waves and moving/static segments; note the discontinuity in speed at the red dashed vertical line, when the Reversal occurs. Black circles indicate the location of the peg contact along the body for sequential frames.

**Figure 6.**
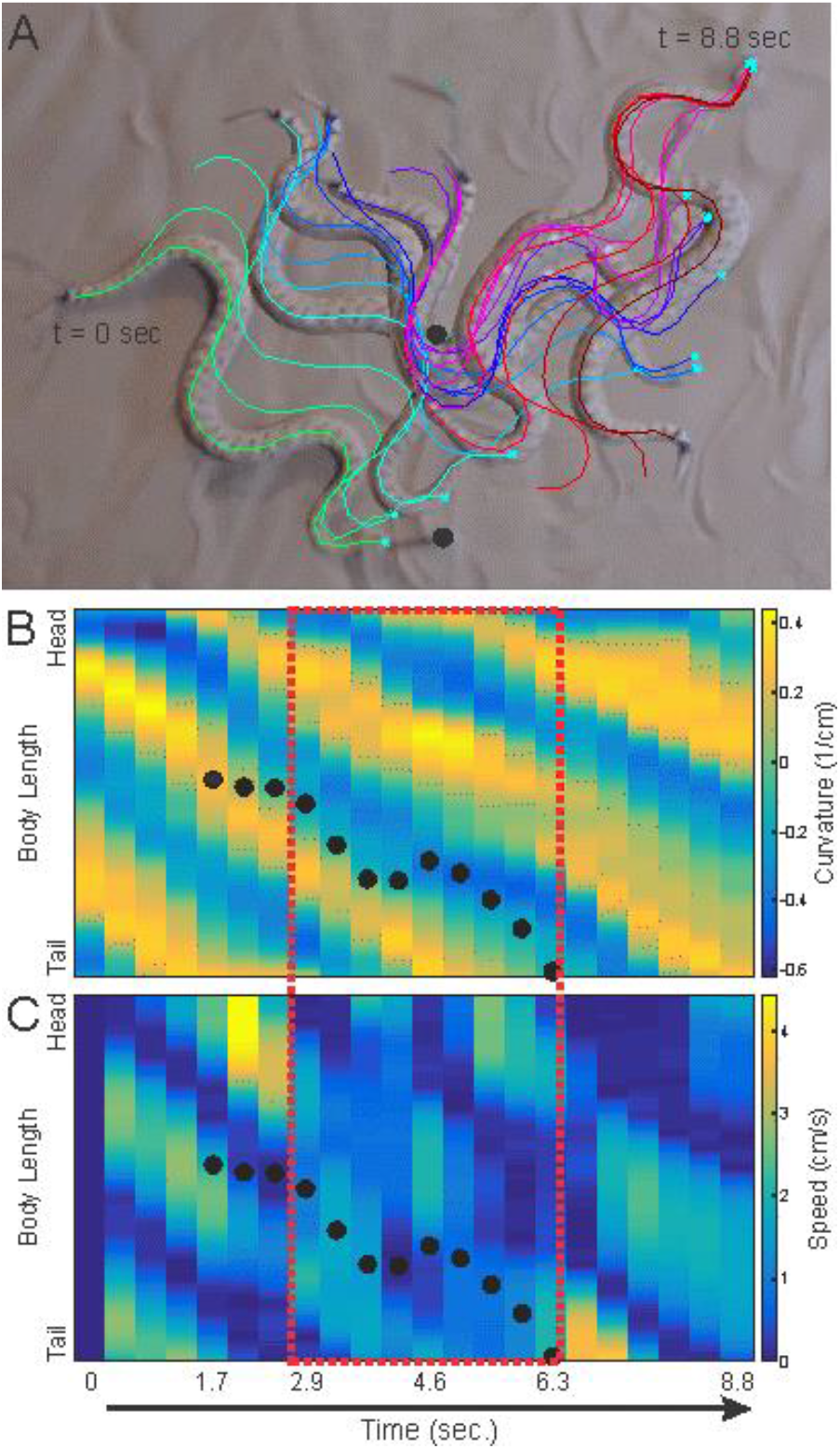
Concertina response. A) Sequential images from Sup. Vid. 9 showing a reversal response, with midline traces at 0.42 second intervals showing the path of the body. The snake is moving from left to right. B) Curvature and C) Speed (derived from midline traces) along the snake’s body over time. The vertical axis is along the length of the body, while the horizontal axis is over time. Bands of curvature and speed going from top left to bottom right show posterior propagation of body waves and moving/static segments; note the transition away from smooth propagation during concertina, shown in the red dashed box. Black circles indicate the location of the peg contact along the body for sequential frames.

Within these categories, snakes would, on rare occasion, change their strategies for the final portion of negotiating the obstacle line. In three runs, sidewinders stopped Propagate Through behaviors to perform concertina locomotion until clear of the pegs, and in another five runs the snakes performed a reversal, at which point it lost contact with the pegs (Fig. 7). Because of the small number of these runs, it is not possible to further analyze what prompted these changes, though in each case they used Propagate Through for the majority of the obstacle negotiation and displayed no obvious differences from the bulk of the runs consisting solely of Propagate Through behaviors. However, when concertina behavior was the initial response, the snake invariably transitioned to either Propagate Through (7/10 runs) or a reversal which allowed the snake to exit the peg line without further contact (3/10 runs) (Fig. 7). When sidewinders performed a reversal turn, the snake was able to pass the peg line without further contact in 17 runs, but otherwise used Propagate Through when contacting a different peg (14 runs) (Fig. 7).In two runs, the initial Reversal led to contact with a different peg, at which time the snake performed a second Reversal and exited the peg line (Fig. 7).

**Figure 7.**
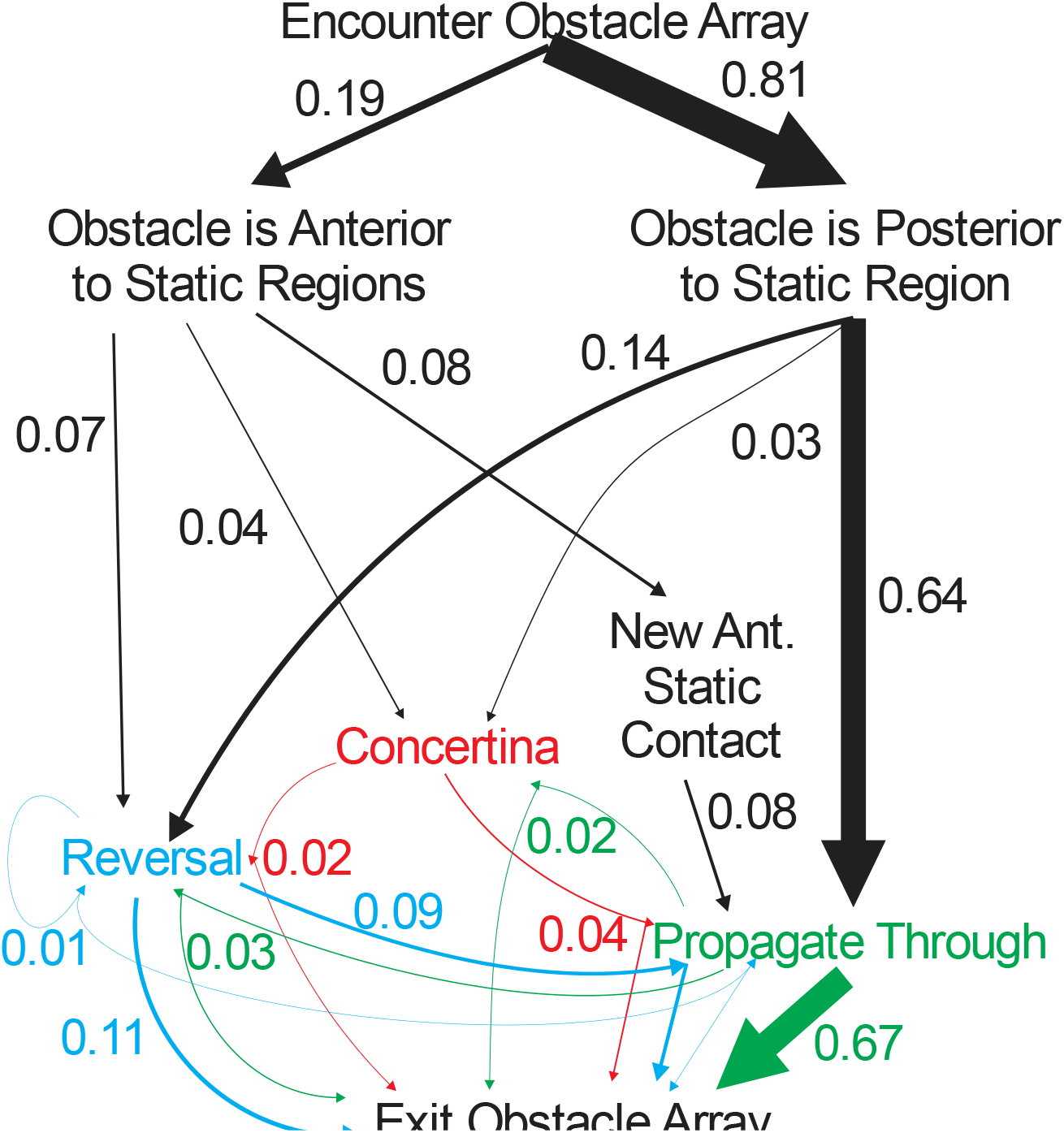
An ethogram (a flow-chart of observed behaviors and transitions between them) of obstacle-traversal behaviors. The number adjacent to each arrow and arrow thickness indicate the proportion of total obstacle encounters involving a given transition between behaviors. The Concertina response is labeled in red, the Propagate Through response is labeled in green, and the Reversal response is labeled in blue.

### Effects of obstacle negotiation

Relative speed of traversal was significantly less than unity for all broad categories of behavior (concertina: mean ± s.d. = 0.359 ± 0.089, t = –22.8, two-tailed p < 0.0001; Propagate Through: 0.539 ± 0.225, t= –22.0, p < 0.0001; Reversal: 0.507 ± 0.154, t=-18.9, p <0.0001), with a significant difference between Propagate Through and concertina values (one-way ANOVA: F2,157 = 3.63, p = 0.0288, Tukey’s HSD q = 3.35), indicating that snakes moved past the peg array at roughly half to a third of their prior speed, a substantial loss in speed (Fig. 8). Snakes also moved more slowly after having cleared the pegs than they did prior to peg contact (9.7 vs. 8.0 cm/s, t = –8.80, two-tailed p < 0.0001) in spite of a slight increase in cycle frequency (0.54 to 0.57 Hz, t = 2.67, two-tailed p = 0.0084), possibly due to a reduction in wavelength (6.7 vs. 6.4 cm, t = –4.30, two-tailed p < 0.0001). No other tested variable showed a significant difference between pre- and post-interaction (Table 1).

**Figure 8.**
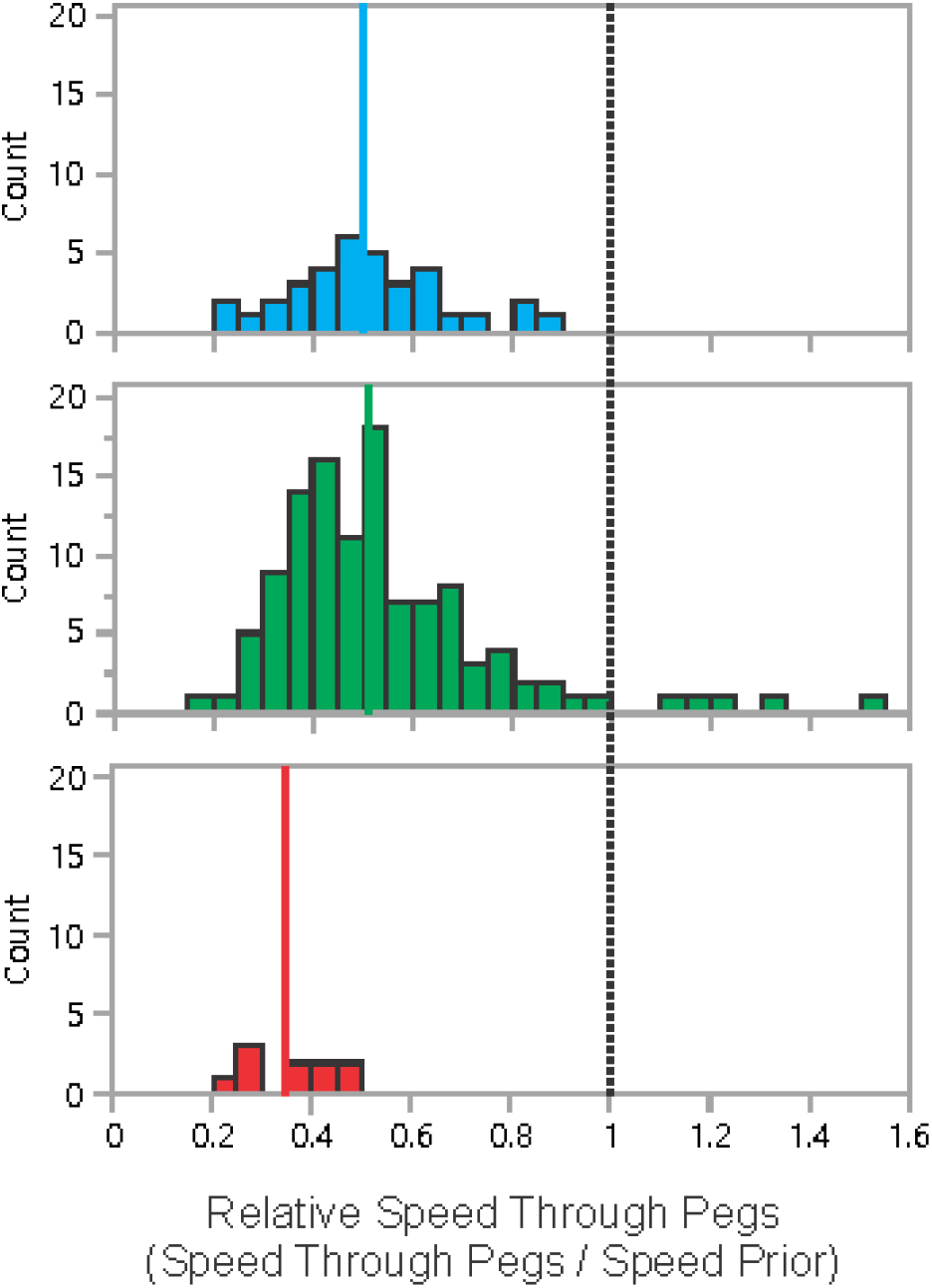
Relative speed of peg traversal. A relative speed of 1 indicates the speed during peg traversal (from initial to final peg contact) was equal to the speed prior to encountering the pegs. Histograms show the distribution of relative speeds for each of the responses, with the median value shown as a solid color line. The Reversal response is labeled in blue, Propagate Through response is labeled in green, and Concertina response is labeled in red.

**Table 1.** Variables predicting behavioral responses.

| Variable | Chi Squared | p | AICc | Relative Likelihood |
| --- | --- | --- | --- | --- |
| Orth. Dist. To Stat. Cont. | 26.07229 | 0.0001 | 219.981 | 1.000000 |
| Contact Point (% BL) | 22.5914 | 0.0001 | 223.462 | 0.175433 |
| Pre-Contact Frequency | 7.526575 | 0.0232 | 238.527 | 0.000094 |
| Pre-Contact Phase | 5.569283 | 0.0618 | 240.484 | 0.000035 |
| Pre-Contact Trajectory Angle | 5.481158 | 0.0645 | 240.572 | 0.000034 |
| Pre-Contact Wavelength | 4.349394 | 0.1136 | 241.704 | 0.000019 |
| Phase At Contact Point | 3.303845 | 0.1917 | 242.749 | 0.000011 |
| Curvature at Contact Point | 3.079601 | 0.2144 | 242.974 | 0.000010 |
| Pre-Contact Speed | 3.05873 | 0.2167 | 242.994 | 0.000010 |
| Number of Static Regions at Time of Contact | 1.759757 | 0.4148 | 244.293 | 0.000005 |
| Pre-Contact Turn Angle | 0.918024 | 0.6319 | 245.135 | 0.000003 |
| Number of Pegs Contacted | 0.232313 | 0.8903 | 245.821 | 0.000002 |

### Effects of peg spacing

The two different peg-spacing distances (10 cm & 15 cm) affected a variety of variables, but there was no effect of peg spacing on the proportion of each response type (Pearson χ^2^ = 2.1, p = 0.35). Runs involving narrower peg spacing had an increased number of pegs contacted (1.32 vs. 1.09, t = −3.58, two-tailed p = 0.0005) and both a lower chance of the first peg contact on the posterior side of the first static contact (Pearson χ^2^ = 18.0, p < 0.0001) and a lower orthogonal distance between the peg and static contact region (1.18 vs. 4.54 cm, t = 4.81, two-tailed p < 0.0001). The most anterior peg impact was more anterior on the body in the narrower peg spacing (0.326 vs. 0.487 body lengths, t = 6.85, two-tailed p < 0.0001). The narrower peg spacing also produced lower relative speed of traversal (0.48 vs 0.58, t = 3.57, two-tailed p = 0.0120) but a higher speed after peg interaction (8.59 vs 7.15 cm/s, t = −2.52, two-tailed p = 0.0127). All wavelength variables showed a slightly higher wavelength for snake encountering the narrower peg spacing, most notably including the wavelength prior to any peg contact (6.90 vs 6.45 cm, t = −2.30, two-tailed p = 0.0033); this suggests that snakes were modulating their waveform prior to contact, potentially to improve outcomes, though wavelength was not a predictor of response (see below).

### Predictors of obstacle negotiation behaviors

Of all the candidate predictor variables, orthogonal distance to static contact showed the most explanatory power (Table 1), with the only other significant variables, point along the body of contact and cycle frequency, showing only a 17% and < 0.0001% chance of offering lower information loss, respectively. Inspection of the raw data showed that the vast majority (102/115) of Propagate Through responses had positive values of orthogonal distance to static contact, compared with only two thirds (23/35) of Reversals and less than half of Concertina responses (4/10) (Fig. 9). Combined with the observations of snake behavior suggesting the crucial role of a static region anterior to the peg contact to provide an “anchor” to pull the posterior portion of the snake’s body through (see above), we examined the impact of reducing the orthogonal distance to static contact to a simple categorical variable coding for whether it was positive or negative (i.e. the peg was posterior or anterior to the first static contact, respectively), and found that there was reduced statistical strength of this model (χ^2^ = 17.70, p < 0.0001, AICc = 228.352, Relative likelihood = 0.0152; Fig. 9). However, snakes could, in the process of propagating through, establish a new static contact region anterior to the prior ones, dramatically changing the position of the contact peg relative to the static contact, switching it from a peg contact anterior to the anterior-most static contact region to posterior to the anterior-most static contact region (Fig. 2BC, Sup.Vid. 4). This modified coding produced the strongest model of all (χ^2^ = 54.08325, p < 0.0001, AICc = 191.97), and by comparison, all other variables showed relative likelihoods of less than 0.00001%. In every case where Propagate Through was the initial response, either the initial peg contact was posterior to a static contact region, or a new static contact region was formed (Fig 7).

**Figure 9.**
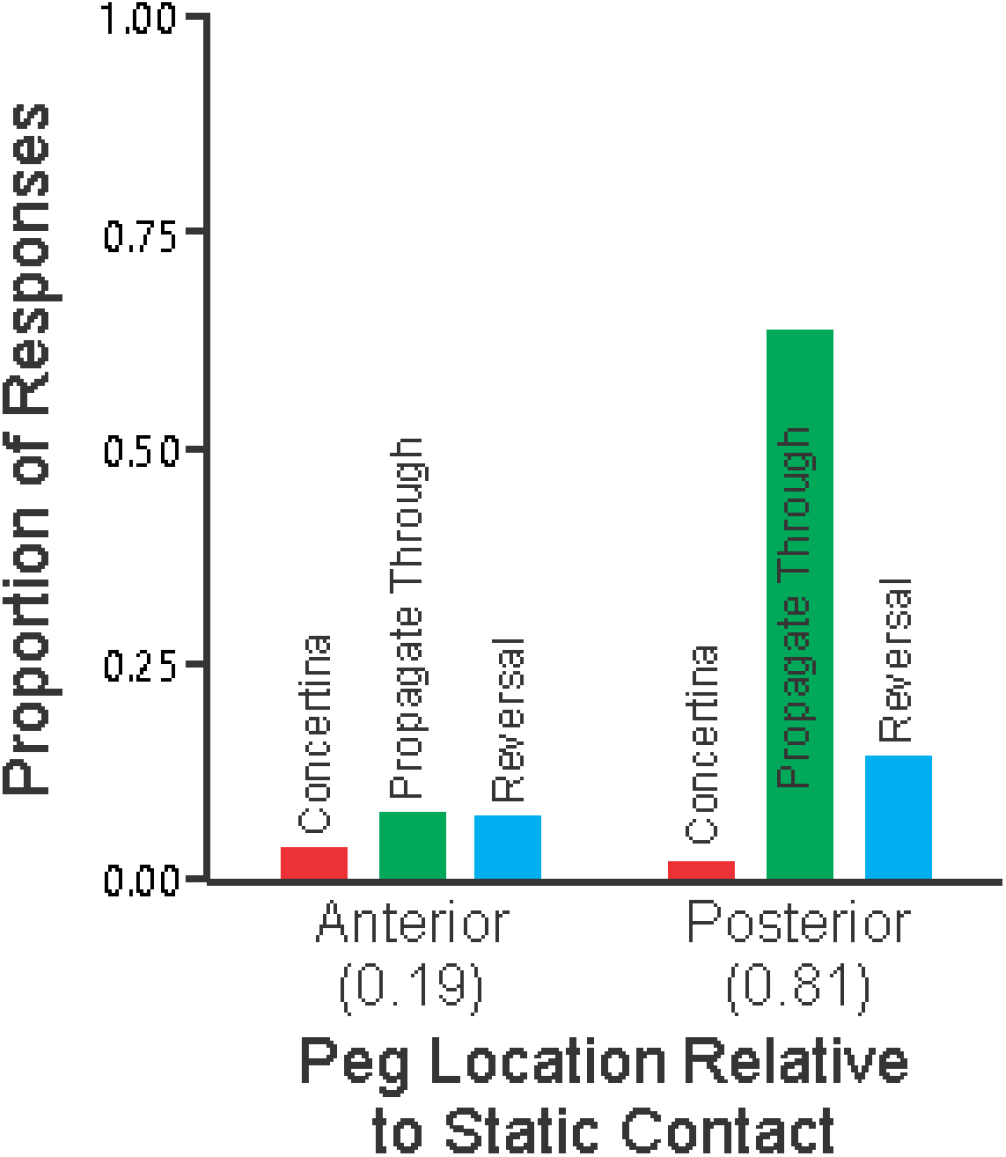
Proportion of responses based on initial peg contact location. The Concertina response is labeled in red, the Propagate Through response is labeled in green, and the Reversal response is labeled in blue.

### Robophysical Testing

Examples of robot performance for the different behaviors are shown in Fig 10 and Supplementary videos 10-12. Trajectories of the robot starting at the same initial position and moving for 5 cycles are shown in Fig 10(i) for (A) nominal behavior on a homogeneous substrate, (B) nominal behavior interacting with a single vertical post, (C) horizontal amplitude increase for one cycle upon contact with post, and (D) spike-improved anchoring plus horizontal amplitude increase upon contact with post. Fig 10(ii) shows the experimentally measured angular positions of the inner six horizontal motors through time. Angular positions vary sinusoidally in time, and waves are initiated at the head and passed down the body. For nominal behavior, every cycle is identical. When contact-sensing information is used to change the behavior, a diagonal band of larger amplitude (dark blue band in Fig. 10(ii)C-D) occurs for one complete cycle. The speed of marked motors along the body is shown in Fig. 10(iii) for all behaviors. When in homogeneous terrain (A), all motors have similar, cyclic speed profiles (Fig 10 A, Sup. Vid 10). In the presence of a post (B), the speed of the posterior segments drops substantially and the head moves faster: the robot’s tail is pinned by the post causing the robot to spin around the post (often never breaking free) (Fig 10 B, Sup. Vid 11). When the amplitude is increased upon contact (C), the tail speed drops and head speed increases (the robot’s head experiences significant sliding), but the increased amplitude allows the robot to squeeze past the post (Fig 10 C, Sup. Vid 12). With the addition of spikes to the second segment of the robot (D), the robot’s head slips less (the speed near the head remains lower throughout the interaction) and the robot is again able to squeeze past the post (Fig 10 D).

**Figure 10.**
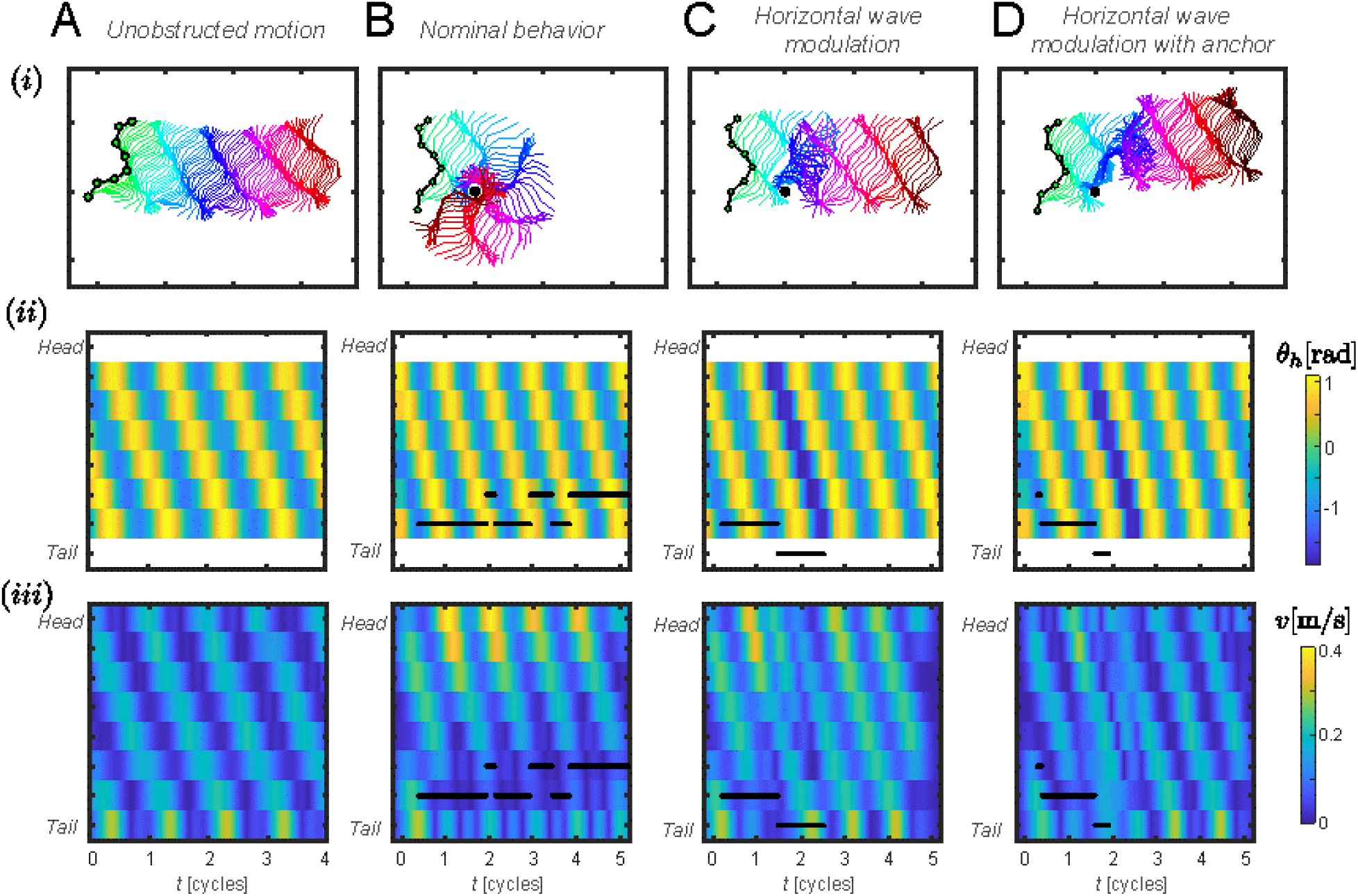
Robot behavioral strategies. (i) Snapshots of robot positions throughout several undulatory cycles, color-coded by time. (A) Robot moving in a homogeneous environment; (B)-(D) Different strategies for moving beyond a single vertical post rigidly affixed to the substrate. (ii) Experimentally measured joint-angles for laterally oriented motors for the strategies shown in (B) no change in behavior upon post contact; (C) at the beginning of the next cycle after contact was sensed, increase lateral amplitude and propagate down the body for one cycle (indicated by the dark blue band); (D) improve anchoring near the head by adding spikes to the second motor and increase lateral amplitude as in (C). (iii) The speed of each lateral joint throughout several cycles.

The effects of initial placement on overall success of the robot in moving beyond the post are shown in Figure 11. The CDFs show that when the robot does not change its behavior upon contacting the post, only about 30% of the trials are able to move beyond the post. Increasing the horizontal amplitude significantly decreases the distribution of pinning times and increases theprobability of success, with the robot moving beyond the post in about 80% of the trials. The addition of spikes to reduce sliding motion near the head further improves performance, yielding smaller pinning times and nearly 100% success rate.

**Figure 11.**
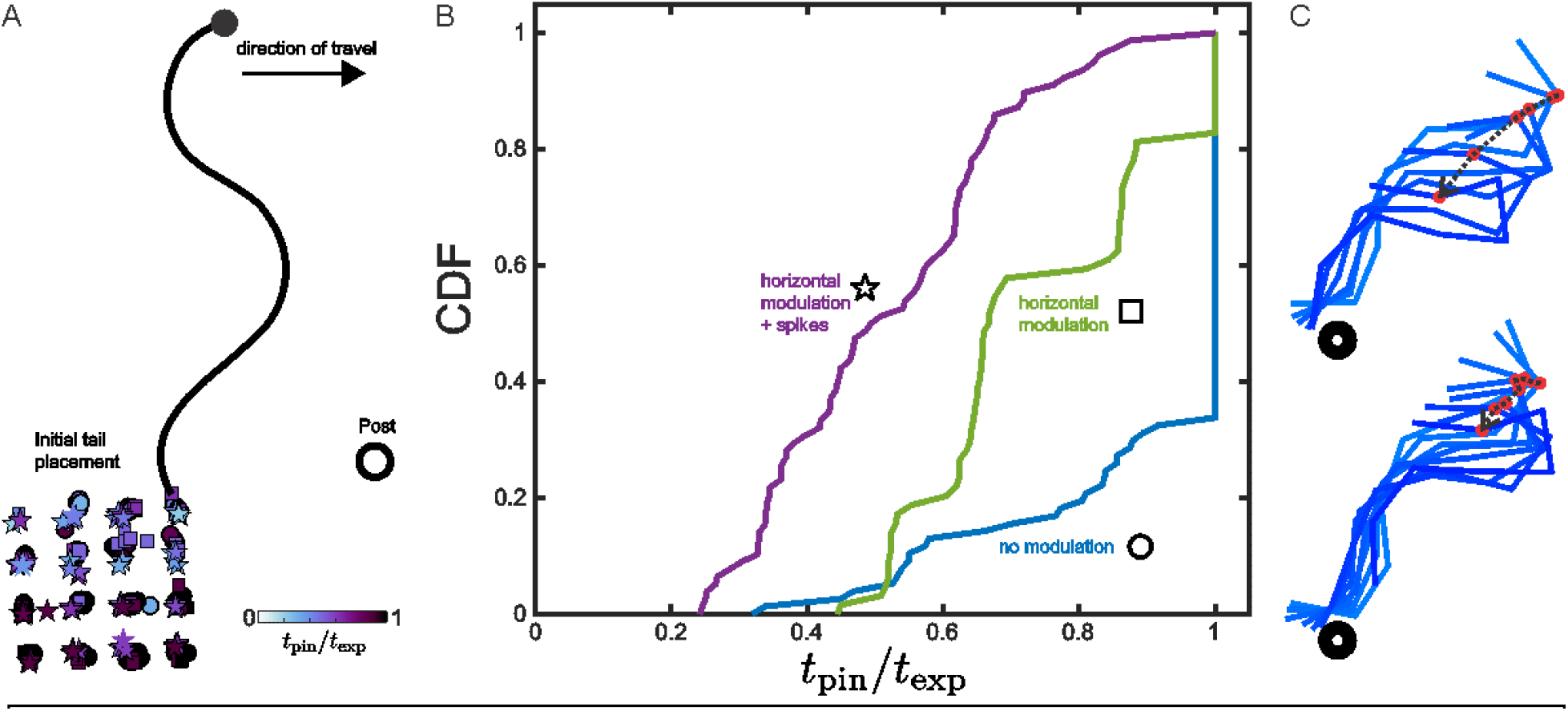
Robot performance across strategies and initial conditions. (A) Initial conditions (defined by robot tail placement) tested for four different control strategies (represented by the different symbol types—see (B) for definitions). The color of each point indicates the estimated duration of each robot–post interaction, normalized by the duration of each experiment (with t = 0) defined at initial contact. (B) CDFs of non-dimensional interaction times for each strategy. t_pin_ / t_exp_ = 0 indicates a short interaction, and t_pin_ / t_exp_ = 1 indicates the interaction lasted the entire experiment. When the robot behavior does not change based on contact, the robot moves beyond the post fewer than 40% of the time. Adding an increase in horizontal modulation greatly improves robot performance, and the robot moves beyond the post in over 80% of the trials. Adding spikes to increase anchoring further improves performance, so that nearly all trials are successful. C) Show the position of Motor 2 (the location of the spikes) during the start of amplitude modulation without (top) and with (bottom) anchoring spikes.

## Discussion

Our results show that, despite the potential to impede or prevent sidewinder, snakes were able to effectively traverse obstacles using sidewinding in almost all trials by modulating their waveform. Furthermore, we were able to replicate the most common waveform modulation, “Propagate Through” in a robophysical model via horizontal wave modulation with contact sensing, demonstrating that this strategy does improve obstacle negotiation behavior and supporting the crucial role of the anterior contact regions in successful obstacle traversal. These results show that, despite the simplicity of the two-wave control template and the potential for obstacles to prevent effective sidewinding, relatively simple modulations of the template can tremendously increase the performance.

The “Propagate Through” response was by far the most common mechanism of moving past obstacles (Fig. 7), as it allowed a continuation of sidewinding with no change in the vertical phase component of the body motions (Astley et al., 2015), and only modulation of the horizontal wave. As the horizontal wave determines the shape and location or ground contact segments, this modulation is likely to cause some slipping or sliding contact. Observations of videos and path-traces (Fig. 4, Sup. Vids. 3-6) show that during “Propagate Through” responses, the posterior ground-contact region shows substantial sliding. However, the anterior ground-contact regions typically maintain static contact throughout, and slipping was only observed briefly in a few instances when the propagating static region intersected directly with the peg and the snake appeared to try to press its body over the obstacle (or deform the obstacle) (Sup. Vid. 6), or the body was pressed against a peg rather than slipping past it (Sup. Vid. 5). During preliminary tests without subsurface anchors, this strategy frequently allowed snakes to knock over obstacles, and may be similarly effective with certain smaller or compliant obstacles in the wild (e.g., dried grass stems). This also suggests that during obstacle-negotiation sidewinding, and possibly during unobstructed sidewinding, the snake is pulling the lifted segment of the body forward via the anterior static contact region to a greater degree than pushing forward from the posterior static contact region. This predominance of pulling from the anterior static contact region may also explain why establishing a static contact region anterior to the obstacle is crucial to implementing the “Propagate Through” behavior. However, the precise distribution of ground reaction-forces during sidewinding remains unknown.

These results were mirrored during robophysical experiments. In the absence of any wave modulation, the sidewinding robot was only able to move past a vertical obstacle approximately 1/3^rd^ of the time (Fig. 11), otherwise it remained simply circling the obstacle (Fig. 10). However, the addition of horizontal wave amplitude modulation greatly improved performance, if at the cost of some slipping of the static contacts, just as was observed in the snake (Figs 4, 10). Similarly, as in the biological snakes, the firm anchoring of the anterior contact region improves peg traversal, as the addition of traction-increasing spikes in the anterior body further improved the performance of the robot (Fig. 11). This serves to both confirm our interpretation of the kinematic data from biological snakes and to highlight to power of modulating the two-wave template to achieve versatile performance while still showing simple control, as previously demonstrated for inclines and turning (Astley et al., 2015; Marvi et al., 2014). Furthermore, these results demonstrate the power of robophysical models for testing biomechanical hypotheses (Aguilar et al., 2016; Astley et al., 2020).

However, while we can replicate the Propagate Through behavior in a robophysical model, it remains unclear what is the exact neuromechanical basis for this behavior, including whether it is centralized or decentralized and whether it requires tactile feedback or is simply a product of the biomechanics of sidewinding. However, previously published muscle-activity data (Jayne, 1988) suggests an intriguing possibility for a passive mechanism (previously shown to improve performance in lateral undulation (Schiebel et al., 2019)) (Fig. 12). For the majority of body segments involved in the lifted phase, muscle activity is unilateral and on the trailing edge of the lifted segment as it moves forward (Jayne, 1988) (Fig. 12). While active force generation can differ from muscle activity (due to activation and deactivation times), this should produce an unopposed torque at those vertebral joints, bending the body towards the trailing edge of the lifted body segment during unobstructed sidewinding (Jayne, 1988) (Fig. 12). This unidirectional force may create a unique response to perturbations, which we term “asymmetrical compliance.” In this proposed model, external perturbations (e.g., impact from an object) opposing the muscle-induced curvature change will be met with increasing resistance from the muscles, either due to reflexes or the intrinsic muscular force-velocity relationship, while external perturbations which complement the actions of the active muscles will be met with minimal resistance (Fig. 12). In the latter case, this mechanism would fall within the broad category of “preflexes”, which provide effectively instantaneous feedback when faced with perturbations (Brown and Loeb, 2000; Full and Koditschek, 1999).

**Figure 12.**
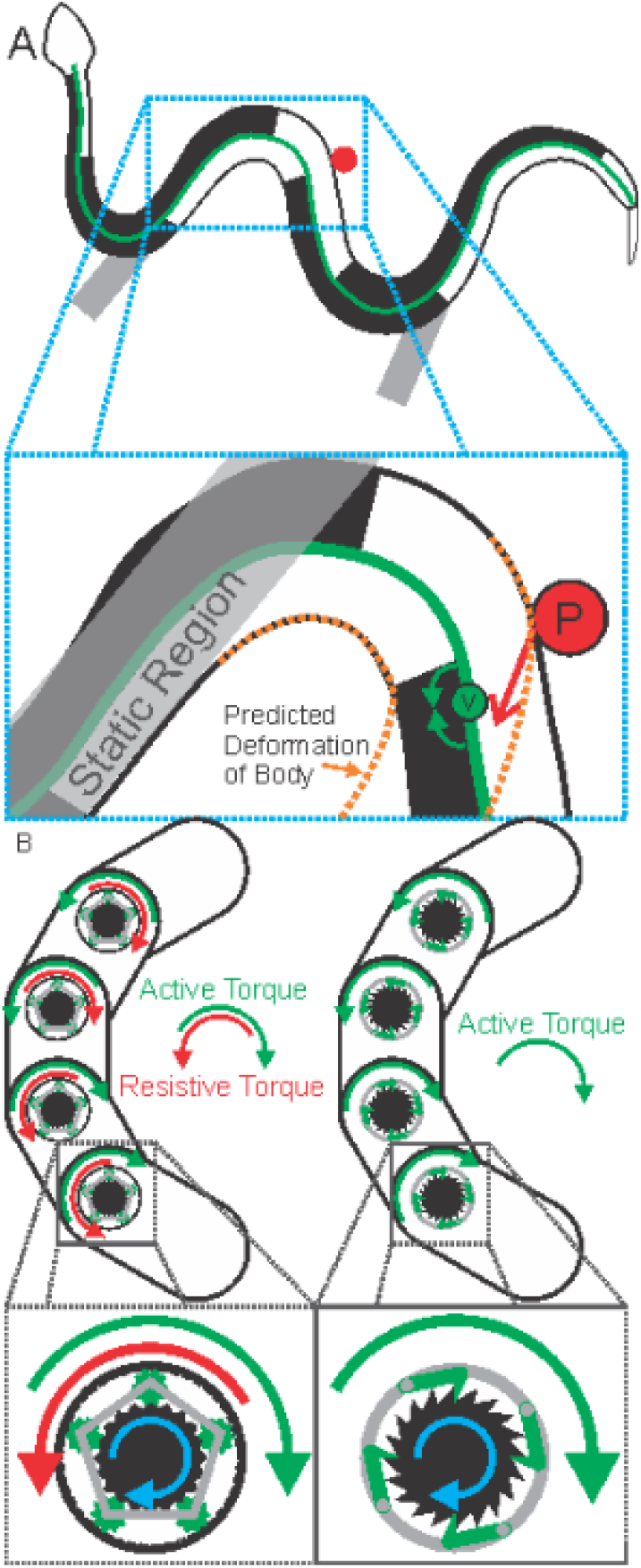
Asymmetrical compliance as a hypothesized mechanism for Propagate Through responses. A) The EMG pattern is based on figures and data from (Jayne, 1988). Black regions indicate electrically active muscle, while white regions are inactive, and the green line is the vertebral column. Grey shading denotes tracks and static contact regions, while the red circle is a peg. The inset figure depicts a hypothesized reaction force from the peg (red arrow), while green circle shows an intervertebral joint and the two curved arrows depict the intervertebral flexion moments due to muscle tension on the trailing side (black). The force exerted by the peg would complement the muscular force, further contributing to flexion at the intervertebral joint. Orange dashed lines show predicted deformation of the body under this control strategy. B) Contrasting diagrams of typical snake robots (left) with a hypothetical alternative for asymmetrical compliance (right). Insets show close-up views of the motors. The standard version uses a gear system (here shown as epicyclic) in which motor torque (blue arrow) applies active torque to deform the body (green arrows), but will resist additional torques in the same direction with a counter-torque (red arrows) due to both the motor and friction of the gear system. The asymmetrical compliance system (here shown with a ratcheting system) will transfer motor torque (blue arrow) to the body (green arrow), but will not resist any external torque in the same direction as the applied torque. These systems are intended to be diagrammatic and not prescribing particular mechanical systems; similar results can be achieved purely in software (Wang et al. 2020).

This “asymmetrical compliance” can be thought of like a torsional spring and ratchet in series, in which externally applied clockwise rotation locks the ratchet and loads the spring, while in counterclockwise rotations the ratchet disengages and the joint spins freely. During sidewinding, the lifted segment is moved through the environment in a direction parallel to the tracks left behind, with the active muscles on the trailing side (Jayne, 1988) (Fig. 12). Thus, at each vertebral joint in the lifted, moving body segment, there is a muscular torque bending it towards the trailing side, but none in the opposing direction. Upon impact with the peg, anterior segments should flex to an even greater degree, while the posterior segments resist bending in the opposite direction, causing this region to “sweep” laterally and drag the posterior static region with it, as we observed in the biological snakes. Furthermore, Jayne’s data showed that at the peak of curvature immediately posterior to the static region had no muscle activity on either side (Jayne, 1988), potentially creating a region of extremely low compliance and localized deformation (contingent on similar lack of activity in other relevant, but yet uncharacterized, muscles). A simulation based on this idea was shown to accurately predict the movement of a laterally undulating snake on sand contacting obstacles such as pegs and walls (Schiebel et al., 2019). Current snake robots typically rely on electrical motors which have very high stiffness in both directions when active, making implementing asymmetrical compliance in a robot difficult (particularly if gear systems add passive stiffness). However, implementing this unusual model of compliance (either via motor control as in (Wang et al., 2020) or alternative actuators) may allow robots to spontaneously manifest adaptive behaviors which currently require sensory feedback and modulation of the motor pattern. Beyond snake locomotion, reciprocal inhibition of antagonistic muscle pairs during activity is a widespread phenomenon (Sherrington, 1905; Tyler and Hutton, 1986), though some co-contraction (at very low antagonist activity) does occur (Baratta et al., 1988), suggesting this principle may have more widespread potential in robot control.

The unique reversal behavior in sidewinders (Astley et al., 2015) offers the snakes a second behavioral option when faced with an obstacle, particularly if the snake’s body configuration and the location of the contact point with the obstacle are incompatible with a “Propagate Through” response (Figs. 5 & 7). Because reversal behavior allows snakes to change their direction of motion without necessarily re-orienting their body, and that change is in direction typically is quite large (Astley et al., 2015), and the obstacle does not impede the use of the turn. This allows the snake to immediately break contact with the initial obstacle, and in some cases, particularly when paired with a subsequent sharp differential turn, the Reversal may allow the snake to escape the obstacles without further contact. However, even when the behavior results in a subsequent impact with another obstacle, the snake’s body configuration and the location of the point of obstacle impact often was compatible with Propagate Through behavior (Fig. 7). In a few rare cases, snakes would repeatedly perform Reversals until Propagate Through became feasible (Sup. Vid. 8). Unlike many other animals which must deal with the obstacles as encountered, sidewinders have a alternative options, allowing additional chances to find a more favorable approach. The use of this alternative behavior suggests an important direction for snake robots research is the development of controllers which can switch between strategies (i.e. amplitude modulation vs reversal) based on local feedback.

These results are broadly compatible with the habitat distribution of sidewinding species, and the rarity of this locomotor mode despite its advantages. Sidewinding is a fast mode of locomotion (Jayne, 1986) with a low cost of transport and high endurance (Secor et al., 1992) enabling longdistance roaming (Secor, 1994; Secor and Nagy, 1994), yet it is only seen in nature in sandy deserts and tidal mudflats with rare and sparse obstacles. Although sidewinding snakes can progress effectively across a wide range of substrates (e.g. sand, smooth & rough wood, tile, cement floors) (Tingle, 2020)(Astley & Mendelson, pers. obs.), this study supports prior suggestions that obstacles impede sidewinding (Mosauer, 1935). Although the snakes were able to effectively negotiate our relatively simple obstacle line, it still significantly impeded their speed and occasionally required direction changes or abandonment of sidewinding in favor of concertina locomotion. Preliminary experiments with a second row of pegs showed similar results only when the pegs were very widely spaced, but total avoidance when even a single row of pegs was spaced more narrowly than trials reported here. While snakes using lateral undulation typically speed up as obstacle density increases (Kelley et al., 1997), we suggest that denser and large obstacle fields will become progressively more difficult to negotiate with sidewinding, and that above a critical threshold it is not possible to move past the obstacles while sidewinding. Thus, we predict that sidewinding will be limited to open spaces, whether sand dunes or mudflats, and in spite of its advantages, sidewinding is unlikely to evolve in more more complex habitats.

## Acknowledgements

We would like to thank Hamid Marvi for the and construction of the test arena, and Zoo Atlanta Keepers David Brothers, Jason Brock, Wade Carruth, and Robert Hill. Kevin and April Young provided considerable assistance with the original collection of the snakes. This research was supported by National Science Foundation (NSF) Grants 1150760 and 0848894; NSF funding for the Student Research Network in the Physics of Living Systems Grant 1205878; Army Research Office Grant W911NF1010343; the Georgia Institute of Technology School of Biology and Elizabeth Smithgall Watts endowment and the Dunn Family Chair Endowment. Research was sponsored by the Army Research Laboratory and was accomplished under Cooperative Agreement No. W9llNF-l0-2-0016. The views and conclusions contained in this document are those of the authors and should not be interpreted as representing the official policies, either expressed or implied, of the Army Research Laboratory or the US Government. The US Government is authorized to reproduce and distribute reprints for Government purposes not withstanding any copyright notation herein.

